# Potent anti-influenza H7 human monoclonal antibody induces separation of hemagglutinin receptor binding head domains

**DOI:** 10.1101/436642

**Authors:** Hannah L. Turner, Jesper Pallesen, Shanshan Lang, Sandhya Bangaru, Sarah Urata, Sheng Li, Christopher A. Cottrell, Charles A. Bowman, James E. Crowe, Ian A. Wilson, Andrew B. Ward

## Abstract

Seasonal influenza virus infections can cause significant morbidity and mortality, but the threat from emergence of a new pandemic influenza strain might have potentially even more devastating consequences. As such, there is intense interest in isolating and characterizing potent neutralizing antibodies that target the hemagglutinin (HA) viral surface glycoprotein. Here, we use cryo-electron microscopy to decipher the mechanism of action of a potent HA head-directed monoclonal antibody bound to an influenza H7 HA. The epitope of the antibody is not solvent accessible in the compact, pre-fusion conformation that typifies all HA structures to date. Instead, the antibody binds between HA head protomers to an epitope that must be partly or transiently exposed in the pre-fusion conformation. The “breathing” of the HA protomers is implied by the exposure of this epitope, which is consistent with metastability of class I fusion proteins. This structure likely therefore represents an early structural intermediate in the viral fusion process. Understanding the extent of transient exposure of conserved neutralizing epitopes also may lead to new opportunities to combat influenza that have not been appreciated previously.

**Author Summary:** A transiently exposed epitope on influenza H7 hemagglutinin represents a new target for neutralizing antibodies.

## Introduction

Influenza displays three glycoproteins that embroider the viral surface: hemagglutinin (HA), neuraminidase (NA) and Matrix-2 ion channel. All of these proteins are necessary for the viral replication cycle. Among these surface glycoproteins, HA is the principal target for neutralizing antibodies. HA is a class I viral fusion protein that facilitates viral entry by interacting with sialic acid receptors on the host cell and then fusing the viral and cell membranes in acidic endosomal compartments. HA is expressed as a precursor form termed HA0, which is cleaved by host cell proteases into HA1 and HA2 domains resulting in a trimer of heterodimers. HA1, the head region, contains the membrane-distal sialic acid receptor-binding site (RBS), while HA2 comprises the stem region and fusion machinery, proximal to the membrane. Proteolytic cleavage at the HA1-HA2 junction liberates the hydrophobic fusion peptide, which then becomes buried in the center of the trimer. The HA1 and HA2 domains remain covalently linked by a disulfide bond after cleavage. This cleaved, pre-fusion conformation is metastable and poised to undergo pH-induced conformational changes but must not do so prematurely.

After influenza virus binds to the host cell, it is endocytosed and trafficked into endosomal compartments, in which the lumen is acidified. Near pH 5.5, HA undergoes large conformational rearrangements, which lead to insertion of its fusion peptide into the host membrane. This process drives fusion of the host and viral membranes and release of the viral RNA genome into the cytoplasm (1).

Influenza evolution, particularly in HA, occurs rapidly with antigenic drift sometimes conferred by single amino-acid changes near the RBS (2). The RBS, which is the most conserved region of the HA head, forms a shallow pocket to which a series of antibodies has been shown to bind and neutralize (3). Although amino-acid mutations in HA are used to evade the host immune system, there are conserved areas of HA that are vital to viral fitness. Head-binding antibodies have been shown to be very effective in neutralizing the virus, but usually only in a strain-specific manner, although some broadly neutralizing antibodies (bnAbs) are known to target the receptor binding site (3,4). Monoclonal antibody (mAb) 5J8, a prototypic head antibody with breadth, targets the RBS via a long HCDR3 that mimics the sialoglycan receptor, and neutralizes H1 strains from 2009 to the pandemic of 1918 (5). RBS antibodies typically have a broader neutralization profile due to sequence conservation of this site, while other regions on the HA1 head are hot spots for strain-specific antibodies (6).

Originating as avian influenza, H7 strains have crossed over intermittently to infect humans (7). Laboratory studies have shown only three amino acids are required to completely change receptor specificity from avian to human, allowing human-to-human transmission and increasing the chance of a new pandemic (8). In fact, an outbreak of H7N9 virus in China in 2013 has been linked to close contact of humans with poultry where chickens, ducks, and pigeons were identified as a reservoir for the virus (7,9). Since then, H7N9 has continued to circulate in poultry reservoirs causing a spike in human infections in recent years (7). Several H7-specific mAbs have been described recently, including H7.137, H7.167, and H7.169, which target highly conserved regions of the HA head adjacent to the RBS (10). Another antibody isolated in the same study, mAb H7.5, was shown to recognize an epitope that overlaps a portion of the H7.167 epitope and neutralize similar strains of human H7 virus, including those from mallards and chickens, but which binds at a different angle (10). The H7.5 mAb potently neutralizes human H7 strains of influenza virus including strains isolated in outbreaks from 2003-2013 in Shanghai, Netherlands, and New York (10).

In the course of our epitope mapping studies of H7-specific mAbs using negative-stain electron microscopy, we noted a striking effect of mAb H7.5 binding to HA in which the soluble H7 ectodomain trimer falls apart. Using biochemical and structural approaches described here, we delineate the phenotypic effect of H7.5 on HA trimers. We also report several cryoEM structures of H7.5 fragment antigen binding (Fab) bound to cleaved or uncleaved H7 HA trimers. Together, these data reveal a previously unappreciated antigenic determinant on HA that, while somewhat inaccessible, may be exploited as a new target for vaccine design given its relative conservation and structural importance. These studies also indicate that the HA trimer is likely sampling subtly different pre-fusion conformations that may provide clues about the early aspects of the conformational changes that accompany the fusion process.

## Results

Antibody H7.5 was described in our previous study that identified several broadly neutralizing antibodies isolated from the peripheral blood B cells of donors who participated in a vaccination trial with monovalent, inactivated influenza (H7N9) A/Shanghai/02/2013 vaccine candidate (DMID13-0033) (10). MAb H7.5 was shown to neutralize H7 viruses and exhibit strong inhibition of hemagglutination activity against a variety of H7 HAs. Preliminary analysis revealed that H7.5 recognizes multiple strains of H7 HAs with K_d_ of binding to HA of less than 0.1 nM, even in the monovalent Fab form, when measured using biolayer interferometry (Fig. S1). The breadth and high affinity for the emerging H7N9 viruses made H7.5 an interesting target for in-depth structural studies. Hence, we first determined the structure of unliganded H7.5 Fab by X-ray crystallography at 2.0 Å resolution (Fig. 1A, Table 1), but its complex with H7 HA could not be obtained despite extensive screening of crystallization conditions.

**Fig. 1.**
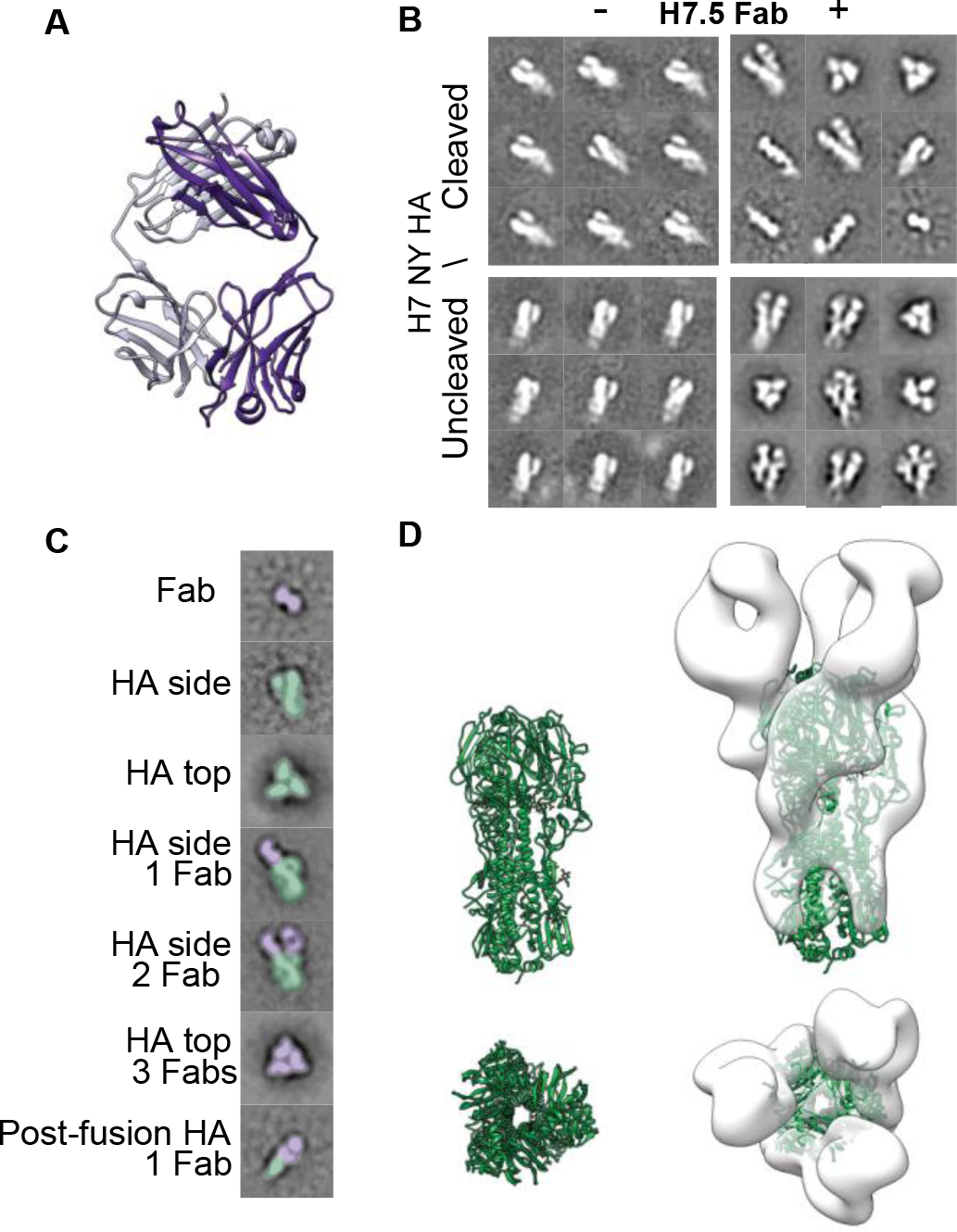
H7.5 Fab crystal structure and nsEM 2D class averages of H7.5 complexed with H7 HA. (A) H7.5 Fab heavy (purple) and light chain (light purple) crystal structure at 2.0 Å resolution. (B) Negative stain EM 2D classes of cleaved and uncleaved H7 NY with and without H7.5 Fab. (C) Example 2D class averages showing range of observations seen after adding H7.5 Fab to H7 HA trimer. The antibody is colored purple and H7 HA colored green. (D) Top and side views of H7 HA1/H2 protomer (green, PDB 3M5G) (left panel), docked into a low-resolution negative stain 3D reconstruction (white surface) of uncleaved H7 NY HA with H7.5 Fab (right panel). Based on the docking, the H7.5-bound HA0 trimer appears to be in a somewhat different conformation than the unliganded HA crystal structure.

**Table 1.** X-ray data collection and refinement statistics for H7.5 Fab.

| **Data collection** | **H7.5 Fab** |
| --- | --- |
| Beamline | APS 23 ID-D |
| Wavelength (Å) | 1.03320 |
| Space group | P22<sub>1</sub>2<sub>1</sub> |
| Unit cell parameters (Å, °) | $a = 65.5$ $b = 99.0$, $c = 208.6$;<br>$\alpha = \beta = \gamma = 90$ |
| Resolution (Å) | 50.00 – 2.00 (2.03 – 2.00) |
| Observations | 624,618 |
| Unique reflections | 91,327 (4,421) |
| Redundancy | 6.8 (6.3) |
| Completeness (%) | 98.5 (96.6) |
| $\langle I/\sigma_I \rangle$ | 21.0 (1.5) |
| $R_{\text{sym}}^a$ | 0.11 (0.74) |
| $R_{\text{pim}}^b$ | 0.05 (0.32) |
| $\text{CC}_{1/2}^c$ | 1.00 (0.80) |
| **Refinement statistics** |  |
| Resolution (Å) | 49.52-2.00 (2.02-2.00) |
| Refs used in refinement | 91,238 (2,795) |
| $R_{\text{work}}$ (%)<sup>d</sup> | 19.9 |
| $R_{\text{free}}$ (%)<sup>e</sup> | 20.2 |
| Protein atoms | 6,555 |
| Waters | 746 |
| Other | 0 |
| **B-value (Å<sup>2</sup>)** |  |
| Average B-value | 36 |
| Protein | 35 |
| Water | 43 |
| Wilson B | 31 |
| **RMSD** |  |
| Bond length (Å) | 0.009 |
| Bond angles (°) | 1.144 |
| **Ramachandran plots (%)<sup>f</sup>** |  |
| Favored | 97.5 |
| Outliers | 0.2 |
| PDB | 6BTJ |

Values in parentheses are for the highest-resolution shell.
<sup>a</sup> $R_{\text{sym}} = \sum_{hkl} \sum_i |I_{hkl,i} - \langle I_{hkl} \rangle| / \sum_{hkl} \sum_i I_{hkl,i}$ and <sup>b</sup> $R_{\text{pim}} = \sum_{hkl} (1/(n-1))^{1/2} \sum_i |I_{hkl,i} - \langle I_{hkl} \rangle| / \sum_{hkl} \sum_i I_{hkl,i}$ , where $I_{hkl,i}$ is the scaled intensity of the $i^{\text{th}}$ measurement of reflection $h, k, l$ , $\langle I_{hkl} \rangle$ is the average intensity for that reflection, and $n$ is the redundancy ( $I$ ).
<sup>c</sup> $\text{CC}_{1/2}$ = Pearson Correlation Coefficient between two random half datasets.
<sup>d</sup> $R_{\text{work}} = \sum_{hkl} |F_o - F_c| / \sum_{hkl} |F_o| \times 100$ .
<sup>e</sup> $R_{\text{free}}$ was calculated as for $R_{\text{work}}$ , but on a test set comprising 5% of the data excluded from refinement.
<sup>f</sup>Calculated using MolProbity (2).

In parallel to the crystallization efforts, we used negative-stain electron microscopy (nsEM) to characterize H7.5 Fab in complex with HA protein from different H7 strains. The difficulty in obtaining crystals of the complex became immediately apparent upon inspection of the complexes in nsEM (Fig. 1B,C). Reference-free 2D classification of cleaved H7 NY HA (A/New York/107/2003) in complex with H7.5 Fab showed substantial heterogeneity relative to previously observed pre-fusion complexes of HA bound to other head-binding antibodies (10) (Fig. 1B,C). In nsEM 2D classes, we observed several different phenomena including variable stoichiometries of bound Fabs and heterogeneous species that were difficult to interpret (Fig. 1B,C). The effect of H7.5 was observed consistently with both cleaved and uncleaved forms of H7 HA from multiple strains including H7 NY HA (A/New York/107/2003), H7 Sh2 HA (A/Shanghai/2/2013) and H7 NL HA (A/Netherlands/219/2003) (Fig. S2). We therefore attempted to capture intermediate conformations of the complex by incubating the cleaved trimer with H7.5 for shorter periods of time. Indeed, a five-minute incubation resulted in enough reliable observations to deduce that H7.5 bound to the HA1 head, but induced an unfamiliar structural phenotype in our 2D classes (Fig. 1B). Within a single data set, all combinations of the H7.5/H7 HA complex stoichiometry were observed (from zero to three Fabs per trimer and individual protomers bound to H7.5 Fab) that probed the conformational landscape of HA (Fig. 1B,C). Relative to other pre-fusion HA structures, the HA1 heads appeared separated, with H7.5 apparently prying them apart or stabilizing more open transient forms of the HA trimer (Fig. 1D).

Unlike cleaved H7 HA trimers that rapidly fell apart into protomers, uncleaved H7 HA (HA0) remained in a trimeric conformation even after overnight incubation with H7.5 (Fig. 1B), although the separation of the HA heads make it distinct from the closed, pre-fusion conformation. This relatively stable complex enabled us to image a larger number of intact particles. Comparison of 2D classes of cleaved and uncleaved trimeric complexes of H7 HA with H7.5 there appears to be no large difference whether zero, one, two or three Fabs are bound (Fig. 1B). Hence, the uncleaved H7 HA, which is stable in the presence of H7.5, is likely to adopt an overall similar structure to the cleaved H7 HA prior to H7.5-induced degradation. We do note that our higher resolution analysis discussed below does indicate there are likely subtle differences in the molecular interactions between protomers of cleaved and uncleaved HA depending on the stoichiometry of H7.5 binding.

Our EM observations demonstrated that H7.5 had a substantial disruptive effect on the pre-fusion structure of HA trimer. To further validate the influence of H7.5 on the HA trimer structure, we investigated the susceptibility of H7 HA to trypsin protease digestion with or without H7.5 bound. Indeed, this experiment showed that addition of H7.5 Fab to H7 Sh2 HA in the presence of trypsin resulted in degradation of HA into many peptidic fragments (Fig. S3). When H7.5 Fab was added to H7 Sh2 HA, 0.1% trypsin was enough to induce the cleavage of H7 Sh2 HA and 2% was sufficient to completely degrade the H7 HA trimer. Without H7.5 Fab, the H7 Sh2 HA trimers remained resistant to protease cleavage up to a trypsin concentration of 2%. These data indicate that H7.5 may prematurely trigger structural changes or fluctuations of H7 HA trimers that acquire protease-sensitive conformations, which is reminiscent of the typical behavior of HA trimers in low pH environments (11,12).

To interrogate the structure of H7.5 bound to H7 HA at higher resolution, we employed single particle cryoEM of the uncleaved H7 NY HA in complex with H7.5 Fab. Reference-free 2D classification resulted in many different views of the complex and secondary structure features were clearly visible (Fig. 2A). The cryoEM 2D classes of the complex in vitreous ice were similar to those observed in nsEM, although the majority of the classes appeared to have three Fabs bound to the HA trimers. Projection images corresponding to meaningful 2D classes were subjected to iterative angular reconstitution and reconstruction. The resulting density map exhibited significant heterogeneity in the Fab densities attached to the head as well as in the membrane-proximal part of the stem. 3D classification was then performed and among the resulting classes, one class was characterized by having well-resolved Fabs bound to the head domain. Data in this class were subjected to further refinement resulting in a three-fold symmetric map with a global resolution of ~3.5 Å (EMDB-9139, PDB 6MLM) that is well resolved in all but the membrane-proximal stem region (Fig. 2B, Table 2). An initial homology model was created using an X-ray structure of the H7 HA1/HA2 protomer (A/New York/30732-1/2005, 3M5G) (13) and combined with our X-ray structure of the H7.5 Fab, and then individually docked into our cryoEM density map exhibiting a nice fit (Fig. 2B, C, D). Next, iterative rebuilding and refinement was carried out in Rosetta (14), yielding a high-resolution atomic model. Since the membrane-proximal part of the map was not well-resolved, a protocol for density-subtracted refinement of this region alone was pursued, resulting in a local density map at ~3.9 Å resolution (Fig. S4). The membrane-proximal part of the model was refined under constraints of this density map and an atomic model was created from combining the two builds. To confirm that local (Brownian) motion was responsible for the disorder in the membrane proximal region – as opposed to the two maps displaying H7 HA in different conformations — coordinates assigned in the density-subtracted local refinement of cryoEM data were applied to the original data and a reconstruction was performed. Indeed, density corresponding to HA1 and to the parts of HA2 not included in local refinement were clearly visible, albeit at an expected lower resolution (Fig. S5), thereby justifying combining the two builds into a meaningful global conformation supported by our experimental observation.

**Fig. 2.**
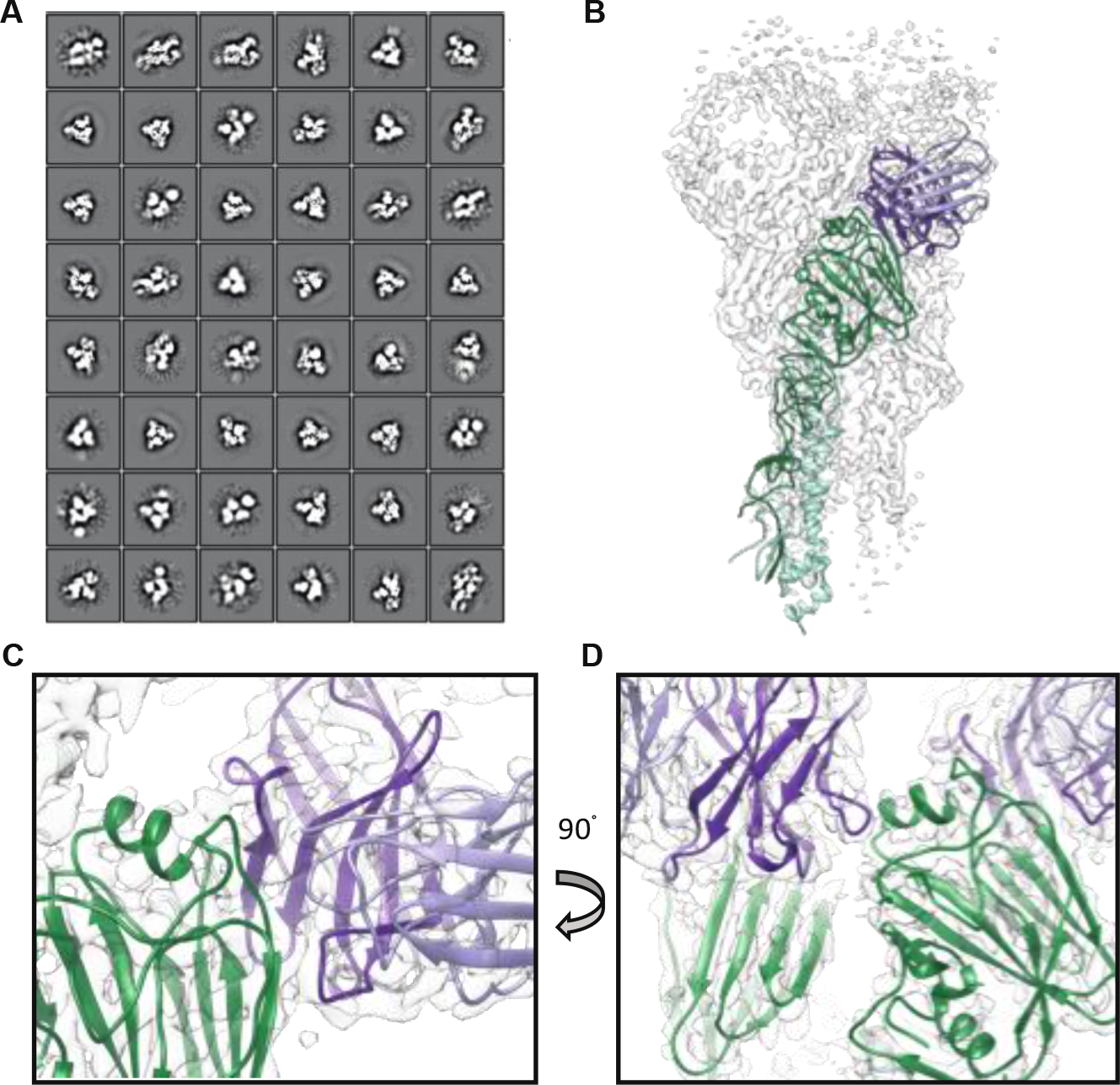
2D and 3D CryoEM image analysis and reconstruction of Fab H7.5 complexed with uncleaved H7 NY HA. (A) 2D class averages produced in Relion of uncleaved H7 NY HA with H7.5 Fab. Multiple views and secondary features are visible. (B) 3D reconstruction (transparent white surface) and H7.5/H7 NY HA atomic model (H7 HA1 in green, HA2 in cyan and H7.5 in light and dark purple). (C) Close up view of H7.5 variable region and HA1 head. (D) Close up view of cross-protomer interaction of H7.5 with neighboring receptor binding domain.

**Table 2.** EM data collection and refinement statistics.

| Map | H7 HA0 | H7 HA0 Stem<br>Base | H7 NL HA1/2- 3<br>H7.5 Fabs | H7 NL HA1/2-2<br>H7.5 Fabs | H7 SH HA1/2-3<br>H7.5 Fabs | H7 NY 1/2-3 H7.5 |
| --- | --- | --- | --- | --- | --- | --- |
| EMDB ID | EMD-9139 | EMD-9139 | EMD-9143 | EMD-9145 | EMD-9142 | EMD-9144 |
| **Data collection** |  |  |  |  |  |  |
| Microscope | FEI Titan Krios | FEI Titan Krios | FEI Talos Arctica | FEI Talos Arctica | FEI Talos Arctica | FEI Technai Spirit |
| Voltage (kV) | 300 | 300 | 200 | 200 | 200 | 120 |
| Detector | Gatan K2 Summit | Gatan K2 Summit | Gatan K2 Summit | Gatan K2 Summit | Gatan K2 Summit | TemCam F416 |
| Recording mode | Counting | Counting | Counting | Counting | Counting | Single Exposure |
| Magnification (incl. post-magnification) | 49,020 | 49,020 | 43,478 | 43,478 | 43,478 | 76,098 |
| Movie micrograph pixelsize (Å) | 1.02 | 1.02 | 1.15 | 1.15 | 1.15 | 2.05 |
| Dose rate (e<sup>-</sup>/[(camera pixel)*s]) | 10 | 10 | 8.375 | 8.375 | 7.576 | 25 |
| Number of frames per movie micrograph | 35 | 35 | 32 | 32 | 32 | 1 |
| Frame exposure time (ms) | 200 | 200 | 250 | 250 | 250 | 500 |
| Movie micrograph exposure time (s) | 7 | 7 | 9 | 9 | 8 | -0.5 |
| Total dose (e<sup>-</sup>/Å<sup>2</sup>) | 67 | 67 | 65 | 65 | 60 | 25 |
| Defocus range (µm) | 0.4-4.0 | 0.4-4.0 | 0.4-5.0 | 0.4-5.0 | 0.5-4.0 | -1.5 |
| **EM data processing** |  |  |  |  |  |  |
| Number of movie micrographs | 1,324 | 1,324 |  |  |  |  |
| Number of molecular projection images in map | 30,032 | 30,032 | 25,146 | 9,682 | 32,117 |  |
| Symmetry | C3 | C3 | C1 | C1 | C1 | C3 |
| Map resolution (FSC 0.143; Å) | 3.5 | 3.9 | 9.2 | 10.2 | 7.4 | 20 |
| Map sharpening B-factor (Å<sup>2</sup>) | -100 | -157 | -259 | -109 | -296 |  |

| Structure Building and Validation |  |  |
| --- | --- | --- |
| PDB ID | 6MLM |  |
| Number of atoms in deposited model | 16710 | 5040 |
| HA1 | 2399 | 517 |
| HA2 | 1370 | 1163 |
| glycans | 42 | 0 |
| MolProbity score | 1.23 (99%) | 1.66 (91%) |
| Clashscore | 2.41 | 4.96 |
| EMRinger score | 2.71 | 1.14 |
| Deviations from ideal, RMSD |  |  |
| Bond length (Å) | 0.013 | 0.015 |
| Bond angles (°) | 1.17 | 1.4 |
| Ramachandran plot |  |  |
| Favored (%) | 96.7 | 94.0 |
| Allowed (%) | 3.0 | 5.0 |
| Outliers (%) | 0.3 | 1.0 |

The epitope on the H7 HA trimer recognized by H7.5 Fab was delineated based on the cryoEM model (Fig. 3A). Intriguingly, the H7.5 antibody was found to simultaneously interact with two separated surfaces on two adjacent HA protomers (Fig. 2D, 3B). Moreover, our model illustrated that the epitope recognized by H7.5 is not completely accessible when all three HA1 heads are close together as observed in the apo cleaved H7 trimer crystal structure. Interestingly, the heavy chain framework region 3 (H-FR3) of H7.5 was juxtaposed to the RBS of an adjacent HA protomer (Fig. S6) and, in a closed trimer model, CDRH3 of H7.5 would clash with residues 189 to 194 in the 190 helix of the adjacent protomer (Fig. 3D).

**Fig. 3.**
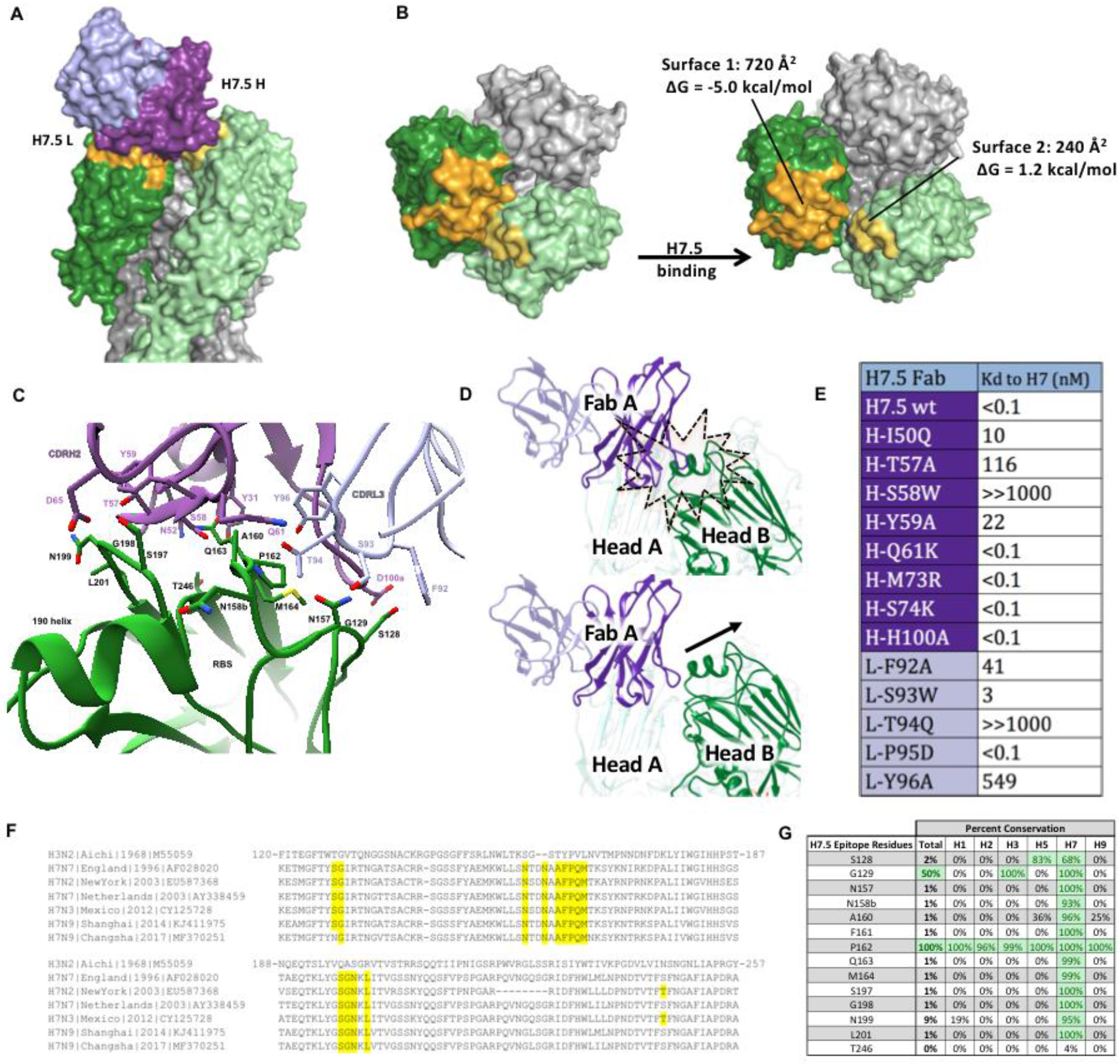
HA protomer movement and mutagenesis. Each H7.5 Fab was found to recognize a primary epitope on one HA1 subunit and a second smaller region on an adjacent HA1 subunit. (A) The variable domain of one H7.5 Fab is shown as a purple surface for the heavy chain and as a light purple surface for the light chain. The HA1 subunit primarily recognized by H7.5 Fab is shown as a dark green surface with the epitope surface color in orange (surface 1). The adjacent subunit that is also contacted by the same Fab was colored in light green, with the epitope region in yellow (surface 2). (B) Top view of the H7.5 epitope on H7 HA before and after H7.5 binding. The estimated buried area surfaces on each HA1 subunit and its predicted Gibbs free energies upon H7.5 binding were analyzed. (C) Interaction between Fab H7.5 and HA primary binding surface 1. The H7.5 antibody is shown as main-chain trace with the interacting residues that are critical to the H7 recognition are shown in sticks. The HA1 subunit with surface 1 is shown in a main-chain trace. The main chain and side chain of H7 HA residues that form direct contacts with H7.5 are shown in stick representation. (D) Interaction between two adjacent protomers and a single Fab. When the H7.5 antibody (purple) is docked onto the closed HA (green) head conformation exhibited in 3M5G, there is a clash with the adjacent protomer (top panel). When the 3M5G HA head is morphed into our model, the increased spacing between the heads alleviates this clash, thereby accommodating the CDRH3 loop (bottom panel). (E) The Kd values of H7.5 or mutated H7.5 Fab binding to the HA1 subunit of H7(A/Shanghai/2/2013) by biolayer interferometry. (F) Sequence alignment of H7 strains with H3N2 reference strain showing the H7.5 epitope (highlighted in yellow) is well conserved among H7. (G) Amino acid sequence analysis of H7.5 epitope among 13,880 influenza A hemagglutinin sequences obtained from www.fludb.org showing poor conservation the H7.5 epitope in subtypes other than H7.

Compared to the apo cleaved H7 trimer model, the buried inter-protomer surfaces in the H7.5 epitope were separated substantially from each other in the H7.5-bound H7 NY HA model (Fig. 3B). The recessed nature of this epitope suggests that one of the surfaces was likely the primary binding interface recognized by H7.5 antibodies, while the other surface allosterically alters upon antibody binding. Indeed, further interface analysis revealed that the HA surface interacting with CDRH2 and CDRL3 of H7.5 (surface 1) generates a total buried surface area of 720 Å^2^, with substantial favorable energy estimated to be -5 kcal/mol and an extensive hydrogen bond network between H7.5 and the HA in surface 1 (Fig. 3B, C). In contrast, the surface between the adjacent HA protomer and H-FR3 of H7.5 (surface 2) is significantly smaller (only 240Å^2^) and appears to be energetically unfavorable with a ΔG of +1.2 kcal/mol (Fig. 3B, S6).

To confirm the structural observations, we performed mutagenesis studies of residues in the antibody paratope by individually reverting the residues to the corresponding germline residues and measuring the affinity changes of the H7.5 mutants to H7 Sh2 HA. Consistent with the interface analysis, mutations in CDRH2 and CDRL3, particularly H-S58W, H-T57A, L-T94Q and L-Y96A of H7.5, drastically reduced binding to H7, while mutations in H-FR3 of H7.5, namely H-M73R and H-S74K, which are proximal to surface 2, did not impact the binding affinity (Fig. 3E, S6). These results strongly support the hypothesis that surface 1 is the primary binding surface driven by favorable energy changes, while surface 2 is a collateral result of the antibody binding event.

Although mutations in H-FR3 or CDRH3 of the H7.5 antibody did not affect binding of the antibody itself, Fab binding requires the adjacent protomer to be displaced from the tight three fold axis at the apex of the trimer to reveal the full epitope (Fig. 3D, S6). The H7.5 epitope therefore includes residues in the inter-HA head contact region (Fig. 3E, S6). Notably, these residues are conserved across all H7 strains from 1996-2017 (Fig. 3F, G). Binding of all three Fabs ultimately results in a conformation of HA with the heads splayed apart.

To further probe the conformational dynamics of the H7 HA when it interacts with H7.5 Fab, we performed hydrogen-deuterium exchange mass spectrometry (HDX-MS) studies on cleaved or un cleaved H7 NL HA in the presence or absence of H7.5 Fab. The differential HDX-MS results of H7 NL HA and H7 NL HA/H7.5 complex were mapped onto the H7 HA Netherlands crystal structure (PDB 4FQV) (Fig. 4A). The HDX-MS data revealed two protected regions (colored in blue) in HA1 of H7 HA upon H7.5 binding. These two discontinuous regions (residues E150 K166, and residues S197-L201) form the potential binding epitope, which is consistent with the cryoEM results (Fig. 3C). Moreover, HDX-MS experiments performed on cleaved H7 NL HA alone indicated a low level of exchange in one of the regions (residue E150-K166), suggesting less solvent accessibility in the binding epitope region. On the other hand, H7.5 binding led to de protection of regions (colored in red) in HA1 and HA2, which indicated that HA became more conformationally flexible to accommodate the binding of multiple H7.5 Fabs.

**Fig. 4.**
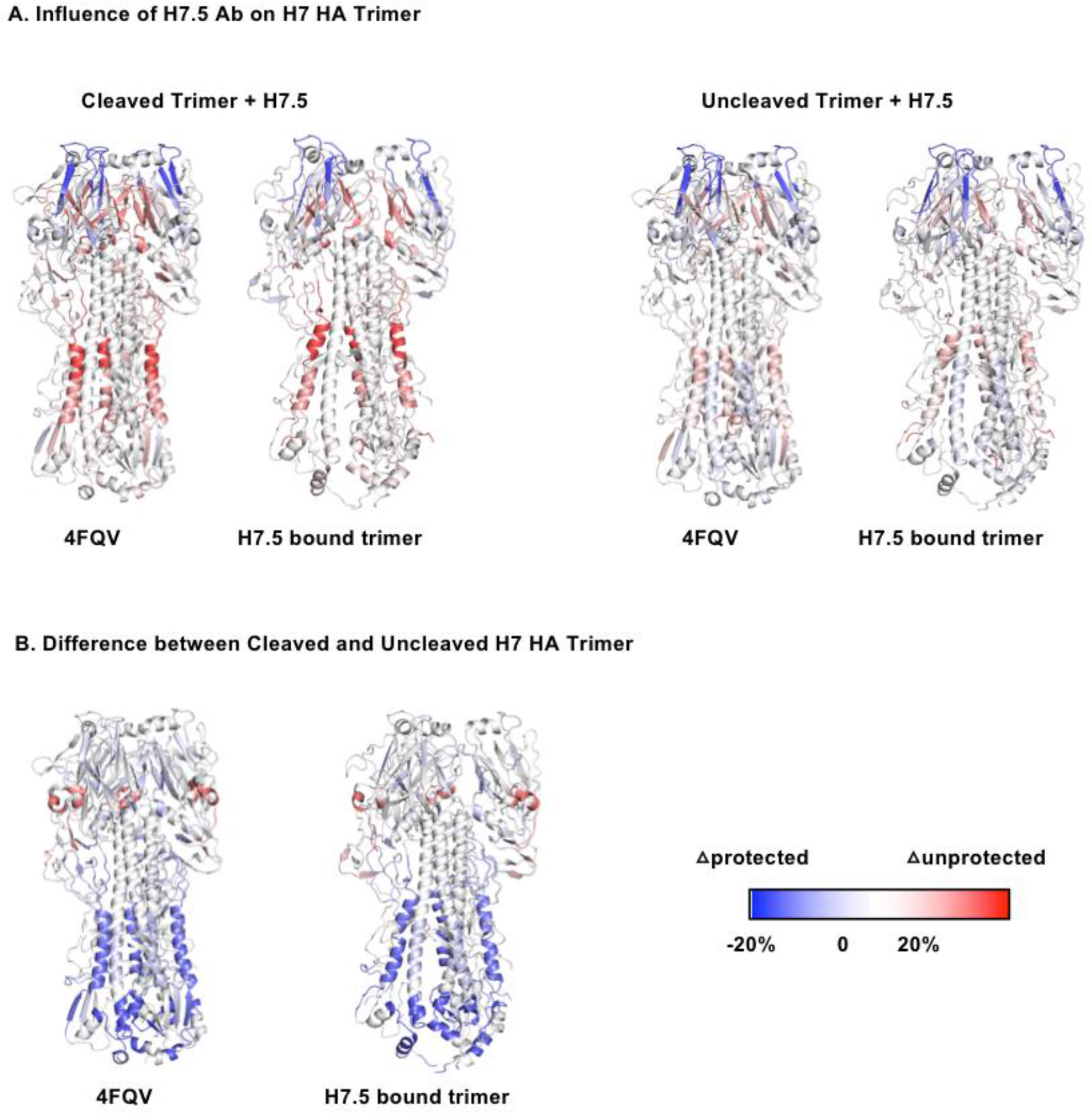
HDX-MS analysis of H7 HA trimer-H7.5 complexes. (A) Difference in HDX profiles of cleaved (left) and uncleaved (right) H7 HA upon H7.5 binding mapped onto closed (4FQV) and H7.5 bound (H7.5/H7 NY HA) model of H7 HA. The epitope of H7.5 in the HA head shows greater protection, consistent with antibody binding. H7.5 binding also induces greater exchange in the HA head region inter-protomer interface, as well as the stem, consistent with structural changes in our models. The effects are stronger in the cleaved HA, consistent with H7.5 induction of trimer dissociation. (B) Regions that are more (blue) or less (red) protected in cleaved H7 HA relative to uncleaved H7 HA are mapped onto closed (4FQV) and H7.5 bound (H7.5/H7 NY HA) model of H7 HA. The stem region of the cleaved HA is more protected from exchange than the uncleaved stem, consistent with maturation of the structure and burying of the fusion peptide in the core of the trimer upon cleavage.

The structural dynamic differences between cleaved and uncleaved H7 NL HA also were compared (Fig. 4B). The HDX-MS results indicated that the cleaved H7 NL HA showed more protection (colored in blue) in the HA2 subunit, compared to uncleaved H7 NL HA, which is more prone to protease cleavage. This finding is consistent with the helical domains in cleaved HA being more stable and is likely related to the maturation of HA from uncleaved HA0 to HA1 and HA2. Detailed results are shown in Figure S7.

To trap additional structural intermediates of HA, we next attempted cryoEM reconstructions of H7.5 Fab bound to cleaved H7 Sh2 HA and H7 NL HA, which differ greatly in season and global location of isolation. Guided by our nsEM experiments (Fig. 1B), we succeeded in capturing an intermediate structure of cleaved H7 HA using a short incubation time with H7.5 Fab immediately prior to sample vitrification for EM experiments. The predominant classes that we observed had three H7.5 Fabs bound to H7 HA and were able to generate asymmetric 3D reconstructions at ~7.4 and ~9.2 Å resolution (EMDB-9142, EMDB-9143) (Fig. 5C, D, S8, Table 2). Similar to the high-resolution structure of H7.5 bound to uncleaved H7 HA, we still observed movement of the HA1 heads away from the apical three-fold axis in the cleaved H7 HAs. Further, in all reconstructions of the cleaved H7 HAs, one of the HA1 heads appeared to be splayed out further than the other two (Fig. 5C, D). In the case of cleaved H7 NL HA, we also refined a class with only two Fabs bound (Fig. S8C).

**Fig. 5.**
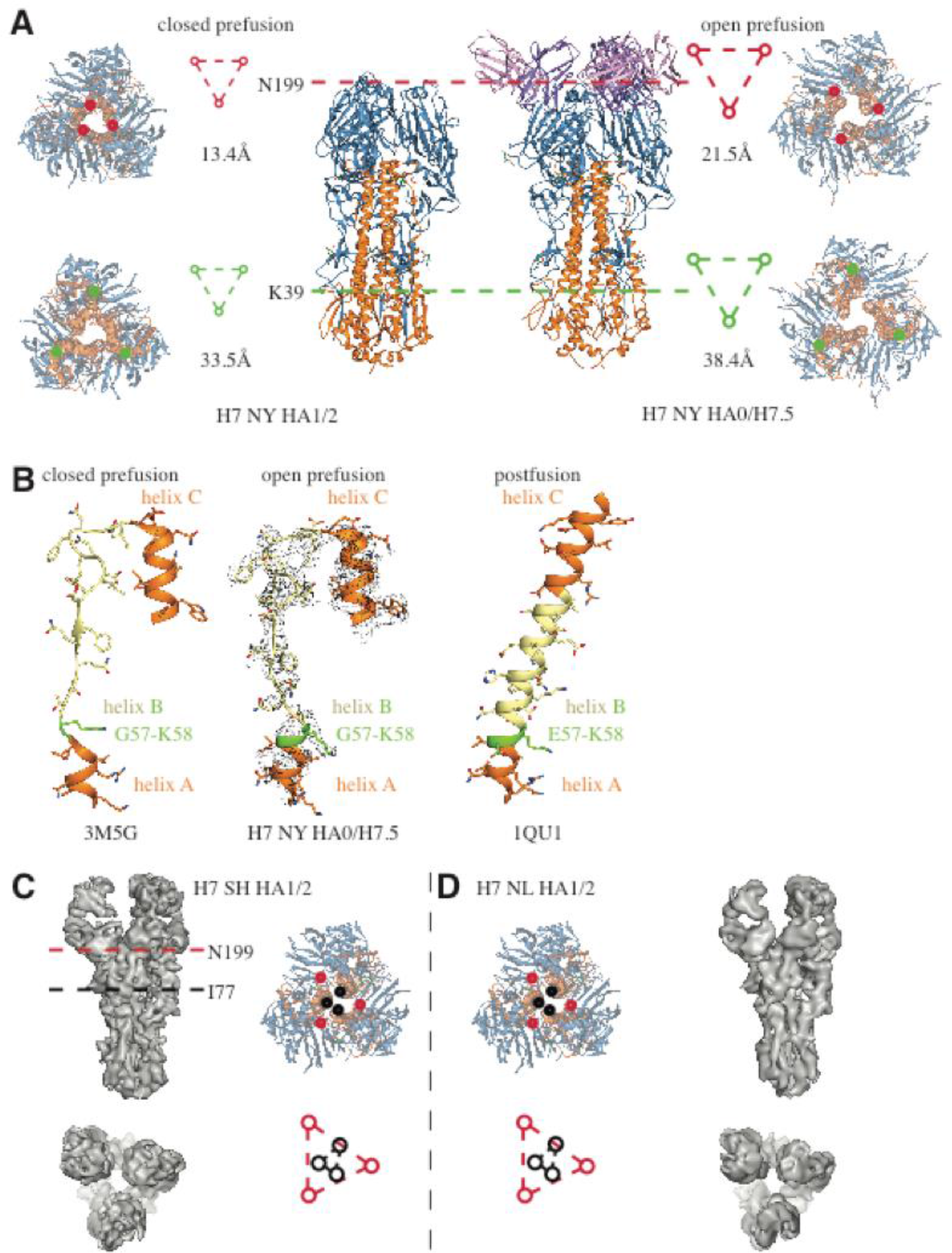
Comparison of H7 strains and movement between unliganded and H7.5 liganded HA trimers. (A) Cross sections of open prefusion trimer (H7.5/H7 NY HA) shows separation in the head domain at N199 (red triangle) and in the stem at K39 (green triangle) when compared to closed prefusion trimer (H7 NY HA) (left side). (B) Residues G57-K58 in HA2 show a transition into a helical conformation after H7.5 binds as seen previously in the post-fusion conformation (PDB 1QU1). Moderate-resolution cryoEM reconstructions of cleaved H7 Sh2 HA1/2 (C) and H7 NL HA1/2 (D). Individual HA1/2 protomers from HA (PDB 3M5G) were fit into the reconstructions to generate pseudo-atomic models. The red and black triangles indicate the spacing of residues N199 (red) and I77 (black) based on the models. Similar to the uncleaved structure in complex with H7.5 shown in (A), the heads move apart relative to prefusion crystal structure. The density maps, reconstructed asymmetrically, appear to have a subtle similar asymmetry, although the limited resolution prevents detailed analysis.

We next compared our high resolution cryoEM derived model of the H7.5 bound to H7 NY HA with separated heads to a crystal structure of HA in the canonical closed pre-fusion conformation (PDB 3M5G). Within a single protomer, our hybrid docking revealed that the HA1 movement was coupled to a subtle change in its position relative to HA2 (Fig. 5). At the trimer level, head separation was accompanied by movement in the HA2 central helices (Fig. 5A). Hence, the HA head motions appear to be communicated downward to the stem, resulting in a small separation of the stem protomers. We hypothesize that this looser packing may trigger events that ultimately liberate the fusion peptide or otherwise destabilize the HA trimer and would lead to initiation of the downhill fusion process, a phenotype that would explain our initial nsEM results with the cleaved HA showing dissociation of the trimer into protomers after incubation with H7.5. Additionally, the head separation correlates with HA2 residues G57-K58 adopting an α-helical conformation (Fig. 5B). These two amino acids were observed previously as a random coil following the C-terminus of HA2 helix A (13). The segment (residues G57-I73) connects helix A with the HA2 central helix; in the stable, trimeric HA2 post-fusion state, this segment was observed to be α-helical forming a continuous extended helix with helix A and the central helix (15). The α-helical organization of residues G57-K58 also correlates with further separation of the membrane-proximal H7 stem. Thus, this structure likely represents an intermediate conformational state of the class I fusion protein of influenza virus. Figure 6 summarizes the results and the events leading up to the trimer dissociation when H7.5 is added.

**Fig 6.**
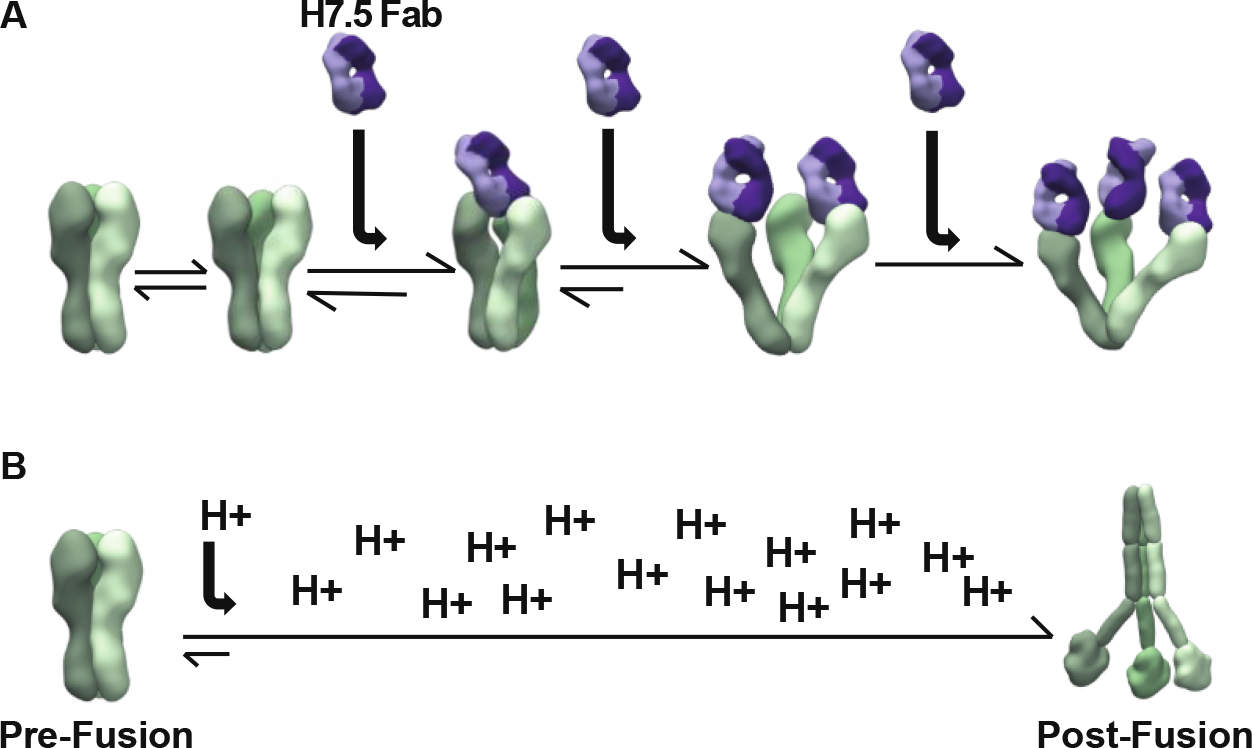
Movement of protomers as antibodies bind and induce the trimer to fall apart. (A) HA trimer heads breathe in equilibrium. When an H7.5 antibody is introduced (purple), it binds to its epitope and prevents re-closing of the HA head, allowing more antibodies to bind. (B) During the fusion process in the endosome, the low pH environment triggers conformational rearrangements in HA.

## Discussion

With the ever-present risk of new avian influenza outbreaks looming, it is important to explore novel ways to target influenza. The recently identified H7.5 recognizes a new conserved epitope on the HA head that can be targeted by neutralizing antibodies. The H7.5 epitope, conserved throughout H7 HA, resides at the HA head protomer-protomer interface and is only transiently accessible. Neutralizing antibodies that bind the HA1 head often target the RBS blocking HA from interacting with sialic acid on host cells (16). Although H7.5 does not bind directly to the RBS, it does however block sialic acid binding to HA (10) and potently blocks intact HA from approaching the cell surface.

The pre-fusion structure of HA has been described extensively, with HA1 and HA2 tightly interacting in a compact arrangement in both cleaved and uncleaved structures (10,17,18). Portions of the post-fusion HA structure have also been crystallized and described (19,20). Interestingly, mAb H7.5 induces premature dissociation without exposure to low pH, and the somewhat open conformation of the receptor binding domains may be reminiscent of early steps in the influenza virus HA-mediated membrane fusion process. After H7.5 binds, it likely induces progressive opening of the trimer and, as more antibodies bind, the trimer is pushed over the activation barrier required for transition into a post-fusion conformation (1). The H7.5 antibody induced decay of HA also represents a new mechanism of influenza neutralization inhibition.

To date, HA has been shown to adopt a stable, closed, pre-fusion conformation (17,21). However, despite significant effort, intermediates in the fusion process have eluded high resolution characterization. It is likely that these intermediates are highly unstable and rapidly transition to the post-fusion form (22). Altogether, our cryoEM studies reveal that HA appears to be somewhat dynamic in its pre-fusion state wherein protomers are able to undergo structural fluctuations. Indeed, we observed movement throughout the HA molecule, including fluctuations in the HA2 stem. The inferred movements of the HA heads to enable antibody H7.5 binding suggest that the HA1 head is “breathing”, and this motion is reminiscent of conformational masking described for the HIV envelope glycoprotein (23-25). Indeed, a very recent study using single molecule FRET observed that H5 HA undergoes reversible conformational changes (26). The structural fluctuations observed in our study, as well as the smFRET study, indicate that movement of the HA1 heads alters interactions between the HA1/HA2 stem portion and the fusion peptide, which reside in the trimer core. This local change then may initiate the large conformational transition toward the post-fusion structure. Whether these movements are similarly coordinated amongst all strains of HA remains to be investigated, but it is likely to be a general property given the metastable nature of the HA fusion machine.

## Materials and Methods

### Preparation of recombinant H7.5 antibodies

H7.5 Fab for crystallization was prepared as previously described(27). In brief, the heavy and light chains of H7.5 were cloned independently into the phCMV3 vector and fused with an N-terminal IgK secretion signal. A His6 tag was added to the C-terminus of the Fab heavy chain. Recombinant cDNAs encoding the Fab heavy and light chains were purified and co-transfected into 293F cells by 293fectin (Invitrogen). After 6-7 days of expression at 37°C, the Fabs were purified from the supernatant by Ni-NTA Superflow (Qiagen) and monoS chromatography (GE Healthcare).

### Crystallization and structure determination of H7.5 Fab

H7.5 Fab was concentrated to 10.0 mg/mL in the buffer of 50mM NaOAc, pH 5.5, for crystallization screening on our high throughput robotic CrystalMation system (Rigaku) at TSRI using sitting-drop vapor diffusion. The best crystals grew in the well with 0.1M MES pH 6.5, 0.01M cobalt chloride, 1.8M ammonium sulfate as mother liquor at 4°C. Crystals were cryoprotected with mother liquor supplemented with 15% ethylene glycol. X-ray diffraction data were collected to 2.00 Å resolution on beamline 23-ID-D at the Advanced Photon Source (APS). The diffraction data were processed with HKL2000 (28) and the structure was determined by molecular replacement with initial models of PDB 3N9G and PDB 4KMT in Phaser (29). Refinements were carried out in Phenix (30), and model rebuilding was performed manually in Coot (31) and the model validated by MolProbity (32).

### Structural analysis

Hydrogen bond and buried molecular surface analyses were calculated using the PDBePISA server of EMBL-EBL. Structure figures were generated by MacPyMOL (DeLano Scientific LLC) and UCSF Chimera package. Chimera is developed by the Resource for Biocomputing, Visualization, and Informatics at the University of California, San Francisco (supported by NIGMS P41-GM103311) (33).

### Preparation of H7 HA

Baculovirus-expressed HA was prepared for the study as previously described(3,34,35). In brief, the HA ectodomain sequence was cloned into the pFastBac vector with an N-terminal gp67 secretion signal peptide, a C-terminal BirA biotinylation site, thrombin cleavage site, foldon trimerization domain, and His6 tag. Recombinant bacmid DNA was generated via the Bac-to-Bac system (Invitrogen) and Baculovirus was generated by transfecting purified bacmid DNA in to Sf9 cells. HAs were expressed by infecting the High Five cells with the recombinant virus, shaking at 110 r.p.m. for 72h at 28 º C. The secreted HAs were purified from the supernatant via Ni-NTA Superflow (Qiagen) and gel filtration (to collect only the HA trimer). The HA trimer was concentrated in the buffer of 20 mM Tris-HCl (pH 8.0) and 150 mM NaCl for storage.

### Trypsin Digestion for Protease Susceptibility

For the protease susceptibility assay, the H7 HA in the experimental group was firstly mixed with H7.5 Fab with molar ratio of 1:3, and diluted to 2 mg/ml (for H7 only) in the buffer of 20 mM Tris-HCl (pH 8.0) and 150 mM NaCl. The control group H7 was diluted with the same buffer to the sample concentration. Both groups of H7 samples were then aloquated and mixed 0, 0.1, 0.2, 1, 2% of trypsin (dissolved in the same buffer above) and each reaction was incubated at room temperature overnight. The samples were then submitted for SDS-PAGE analysis to determine the stability of the H7 protein.

### Bio-Layer Inferometry

An Octet RED instrument (FortéBio, Inc.) was used to determine K_d_ of the antibody-antigen interactions by bio-layer interferometry. To determine the binding affinity of H7.5 Fab to its ligands, HAs (biotinylated as previously described (3,34) were immobilized onto streptavidin coated biosensors (FortéBio, Inc.) and incubated with H7.5 Fab at 62.5-500 nM for 120s for association and then incubated in PBS buffer with 0.2% BSA for dissociate for 120s. The signals for each binding events were measured in real-time and K_d_ values were calculated by fitting to a 1:1 bivalent analyte model. In this case, the Kd values were estimated to be less than 10^-3^ nM, as no dissociated was observed.

### Negative stain electron microscopy

H7.5 IgG was digested with 4% papain w/w for 4 hours before adding iodoacetamide. The Fab was purified using a protein A column and then added to H7 NY HA with a 3x molar excess of Fab and incubated for 30 minutes at room temperature. The complex was purified using an S200i column (GE Healthcare, Amersham, UK) and immediately added to 400 mesh copper grids with 2% uranyl formate. Micrographs were taken on a Tecnai Spirit with a TemCam F416 camera using Leginon (36). Particles were selected using DoGpicker (37) and then stacked and aligned into 2D classes in Appion (38) with MRA/MSA (39). Particles not representing HA trimers or the complex were removed and a clean stack was processed in RELION (40).

### Cryo electron microscopy

Sample preparation for uncleaved H7 HA was similar to negative stain. Directly after SEC purification, the complex was added at 1mg/mL to 2/2 gold Quantifoil grids with amphipol and frozen in liquid ethane using a Vitrobot. Cryo sample preparation for cleaved H7 HA with H7.5 Fab does not include a column purification step. Complexes were formed, incubated for one minute at room temp and immediately frozen.

### High resolution image processing

1,324 micrographs were recorded of the uncleaved H7 NY HA/H7.5 Fab complex on a Gatan K2 summit detector mounted on a Titan Krios operating at 300 kV. Data were collected in counting mode at a nominal magnification of 29,000x. Dose rate was ~10 electrons/(camera_pixel*s) and frame exposure time was 200 ms. Total exposure time was 10s with a total dose of 67 electrons/Å^2^. Movie micrograph frames were aligned using MotionCor2 (41), dose-weighted and integrated. CTF models were determined in GCTF (42). Projection image candidates of H7 NY HA/H7.5 were identified using a difference-of-Gaussians picker (37). The resulting set of candidate projection images were subjected to 2D class averaging implemented in RELION 2.1b1 (40). 227,202 projection images that generated well formed class averages were selected for further data processing. Iterative angular reconstitution and reconstruction was attempted. Due to on-symmetry axis preferred orientations, reconstruction artefacts were insurmountable for asymmetric refinement. C3 symmetry was imposed in a second attempt and a stable, albeit somewhat stretched, 3D reconstruction of the data set was obtained. The data set was then subjected to 3D classification and a subclass of the data (30,032 projection images) was identified; its reconstruction was characterized by persistent Fab densities — indicating full stoichiometric occupancy — and no obvious reconstruction artefacts. This subclass of data was then refined under C3 symmetry constraint to a final resolution of 3.5 Å. Fab Fv regions were well-resolved as were HA1 and part of HA2. The membrane-proximal stem of HA2 was largely disordered. Attempting to recover a structure of the membrane-proximal stem, a globular mask was placed encompassing the disordered part of the map. Inverting the mask, the ordered part of the map could be isolated from which a mask was created to isolate the constant part of the original map. This constant part map was then used to project onto R2 along projection directions deduced from Euler and X,Y coordinate assignments obtained during the earlier-referenced refinement, followed by subtraction from the original projection images. This density-subtracted data set was subjected to further iterative angular reconstitution and reconstruction resulting in a density map of the membrane-proximal stem resolved to 3.9 Å. To investigate if the two maps could be combined and interpreted as one instance of multiple only locally diverging — breathing — conformations, the Euler and X,Y coordinates determined in the stem-refinement was applied to the corresponding original projection images and the coinciding 3D object reconstructed. This density map exhibited, albeit noisy, density for the constant part of the structure suggesting that the membrane-proximal part of the stem is breathing locally around the position determined in the local map. Treatment of our cleaved H7 NL HA/H7.5 and H7 Sh2 HA/H7.5 datasets was performed in RELION 2.1b1 and followed a standard procedure as outlined above prior to local refinement procedures and as outlined in the supplemental information (Fig. S5). FSC 0.143 between independently refined half sets were 7.4 Å (H7 Sh2 HA/3 H7.5 Fabs), 9.2 Å (H7 NL HA/3 H7.5 Fabs) and 10.2 Å (H7 NL HA/2 H7.5 Fabs).

### Model building and refinement

A homology model was created (Modeller; (43)) based on the X-ray structure of HA7 HA1/H2 (A/New York/30732-1/2005; PDB 3M5G) (13). The model was created by combining with the H7.5 Fab X-ray structure and rigid-body fitted to the cryoEM density map. A fragment library consisting of 200 7mers for each amino acid position in the model was compiled from a non-redundant protein structure database. Iterations of manual and Rosetta fragment-library-based centroid rebuilding-and-refinement was then performed (14). The resulting model was all-atom-refined under constraints of the density map. Glycans were manually built in COOT (31) and final rounds of real-space refinement performed in PHENIX 1.12(44). The membrane-proximal part of HA2 was built similarly in the local map referenced above. The builds were then combined into the final structure. Evaluation of builds were performed in MolProbity(32), EMRinger(45) and Privateer(46) and by the PDB validation server.

### Hydrogen deuterium exchange Mass Spectrometry (HDX-MS)

Antigen-Antibody complexes were prepared by mixing HA H7 and H7.5 antibodies at 1:1.1 stoichiometric ratio, and incubating at room temperature for 30 minutes, and then kept at 0 °C. For control experiments, free antigens were also diluted with same volume of protein buffer and treated same way as complexes. To initiate hydrogen-deuterium exchange reactions, 2 μl of pre-chilled protein stock solution (free un-cleaved HA NL H7, 1.8 mg/ml; H7.5-uncleaved HA NL H7, 4.5 mg/ml, cleaved HA NL H7, 1.6 mg/ml, or H7.5-cleaved HA NL H7, 4.5 mg/ml) was diluted into 4 μl D_2_O buffer (8.3 mM Tris, 150 mM NaCl, in D_2_O, pDREAD 7.2) at 0 °C. At the indicated times of 10 sec, 100 sec, 1000 sec, 10000 sec and 100000 sec, the exchange reaction was quenched by the addition of 9 μl of optimized quench solution (6.4M GuHCl, 1 M TCEP, 0.8% formic acid, pH 2.4) at 0 °C. After incubating on ice for 5 min, the quenched sample was diluted 5 fold with 0.8% formic acid containing 16.6% glycerol, immediately frozen on dry ice and stored at -80°C. In addition, un deuterated samples and equilibrium-deuterated control samples were also prepared. All samples were then loaded onto our in house DXMS apparatus for online digestion and separation. The resulting peptides were directed into an OrbiTrap Elite Mass Spectrometer (Thermo Fisher Scientific, San Jose, CA) for DXMS analysis. Instrument settings have been optimized for HDX analysis. The data acquisition was carried out in a data-dependent mode and the five or ten most abundant ions were selected for MS/MS analysis. Proteome Discoverer software was used for peptide identification. The centroids of each peptide was calculated with HDExaminer, and then converted to corresponding deuteration levels with corrections for back-exchange. The deuteration levels of the peptides were further sublocalized using overlapping peptides by MATLab program.

### Epitope Conservation Analysis

Non-duplicate hemagglutinin sequences were fetched from the databases at www.fludb.org in FASTA format. Strain A/Aichi/2/1968 HA gene (Genbank Accession M55059) was then defined as a reference for residue numbering. We then used an in house script to build a database of every HA sequence’s residues at every position relative to the reference. First, we performed pairwise Clustal alignment of every sequence to the reference sequence (47). The results were parsed into a database keyed on positions relative to the reference, with gaps notated as sub-positions following the last identical residue. We are then able to interrogate the database by providing position numbers and desired residues at the position, providing an output of sequences that meet the search criteria, as well as breakdowns of residues at the positions.

## Data Availability

The final coordinates for the H7.5 Fab structure have been deposited to the RCSB database with the accession code PDB 6BTJ. The nsEM map of H7 NY HA1/2 complexed with 3 H7.5 Fabs bound, and the cryo-EM maps of H7 NL HA1/2 with 3 H7.5 Fabs bound, the H7 NL HA1/2 with 2 H7.5 Fabs bound and the H7 Sh2 HA1/2 with 3 H7.5 Fabs bound were deposited to the Electron Microscopy Databank with the accession codes EMD-9144, EMD-9143, EMD-9145 EMD-9142 respectively. The cryo-EM map and fitted coordinates for H7 NY HA with 3 H7.7 Fabs bound has been deposited to the RCSB database with accession numbers EMD-9139/PDB 6MLM.

## Author contributions

EM data collection and processing: H.L.T, J.P. and A.B.W. EM model building and validation: H.L.T., J.P., C.A.C., and A.B.W. HA sequence analysis: C.A.C. and C.A.B. X-ray crystallography data collection and processing: S.L. and I.A.W. Protein production: S.L. and S.B. HDX-MS: S.U. and S.Li. Manuscript writing and revision: H.L.T., J.P., S.L., S.B., C.A.C., C.A.B., I.A.W., J.E.C., and A.B.W.

## Acknowledgments

We thank Robert Kirchdoerfer for help with data processing and interpretation, Bill Anderson and J.C. Ducom for assistance with microscope management and computational support, Lauren Holden for critical reading of the manuscript. This work was supported by NIH grant U19 AI117905 and NIH contract HHSN272201400024C. This research used resources of the Advanced Photon Source, a U.S. Department of Energy (DOE) Office of Science User Facility operated for the DOE Office of Science by Argonne National Laboratory under Contract No. DE-AC02-06CH11357. The contents of this publication are solely the responsibility of the authors and do not necessarily represent the official views of NIAID, DOE or NIH.

